# Aging decreases the lateralization of gravity-related effort minimization during vertical arm movements

**DOI:** 10.1101/2021.10.26.465988

**Authors:** Gabriel Poirier, Charalambos Papaxanthis, Mélanie Lebigre, Adrien Juranville, Robin Mathieu, Timothée Savoye-Laurens, Patrick Manckoundia, France Mourey, Jeremie Gaveau

**Author notes:** Corresponding author: Gabriel Poirier.

## Abstract

Motor lateralization refers to differences in the neural organization of cerebral hemispheres, resulting in different control specializations between the dominant and the non-dominant motor systems. Multiple studies proposed that the dominant hemisphere is specialized for open-loop optimization-like processes. Recently, comparing arm kinematics between upward and downward movements, we found that the dominant arm outperformed the non-dominant one regarding gravity-related motor optimization in healthy young participants. The literature about aging effects on motor control presents several neurophysiological and behavioral evidences for an age-related reduction of motor lateralization. Here, we compare the lateralization of a well-known gravity-related optimal motor control process between young and older adults. Forty-one healthy young (mean age = 24.3 ± 3 years) and forty-two healthy older adults (mean age = 72.0 ± 6 years) performed single degree-of-freedom vertical arm movements between two targets (upward and downward). Participants alternatively reached with their dominant and non-dominant arms. We recorded arm kinematics and electromyographic activities of the prime movers (Anterior and Posterior Deltoids) and we analyzed parameters thought to represent the hallmark of the gravity-related optimization process (i.e directional asymmetries and negative epochs on the phasic EMG activity). We found strong age x arm interaction effects on all parameters; i.e., relative durations to peak acceleration and peak velocity and the negativity of antigravity muscles’ phasic signals. Although all three parameters showed a lateralization effect where the dominant arm was superior to the non-dominant arm in young adults (as in Poirier et al. 2022), we found no such effect in older adults. With both arms, the results of older adults lied between those of the dominant and non-dominant arm of young adults. These results add to those of recent literature showing that feedforward motor control remains functional in older adults. More, the results obtained with the non-dominant arm may support a previously hypothesized increased reliance on predictive mechanisms in older adults.

## Introduction

Aging causes several behavioral modifications. Notably, age-related brain alterations are known to affect motor control processes (Seidler et al., 2010; Papegaaij et al., 2014; Poirier et al., 2021). Studying behavioral modifications is important to improve the understanding of aging effects on the neural implementation of behavior and, thus, the prevention and rehabilitation of age-related loss of mobility (Krakauer et al., 2017; Poirier et al., 2021).

An important notion in motor control is the concept of motor lateralization. This concept refers to the differences in the neural organization of the cerebral hemispheres, resulting in different specializations between dominant and non-dominant motor systems regarding motor control (Flowers, 1975; Roy and Elliott, 1986; Carson et al., 1992; Roy et al., 1994; Bagesteiro and Sainburg, 2002; Sainburg, 2002, 2014; Woytowicz et al., 2018; Jayasinghe et al., 2021). Multiple studies proposed that the non-dominant hemisphere is specialized for closed-loop impedance control that stabilized arm postures against unexpected environmental conditions, whereas the dominant hemisphere would be specialized for open-loop optimization-like processes (Bagesteiro and Sainburg, 2002; Sainburg, 2002; Yadav and Sainburg, 2011, 2014; Schaffer and Sainburg, 2017). For example, the results of Schaffer and Sainburg, (2017) suggest that the dominant arm better takes advantage of gravity during unsupported reaching movements, compared to the non-dominant arm. It must be noted, however, that some recent results may limit this theory (Takagi et al., 2020; Maurus et al., 2021).

We recently investigated inter-limb differences regarding gravity-related optimization processes during vertical arm movements (Poirier et al., 2022). A large body of literature combining mathematical simulations and behavioral experiments previously demonstrated that the brain optimally takes advantage of gravity torque to minimize muscle effort during vertical arm movements (Berret et al., 2008; Crevecoeur et al., 2009; Gaveau et al., 2011, 2014, 2016, 2021). More precisely, investigating vertical arm movements, multiple studies have reported direction-dependent kinematics where upward movements exhibit sharper acceleration profiles than downward ones (Papaxanthis et al., 2005; Gentili et al., 2007; Le Seac’h and McIntyre, 2007; Gaveau and Papaxanthis, 2011b; Gaveau et al., 2011, 2014, 2016, 2021; Yamamoto and Kushiro, 2014; Yamamoto et al., 2016, 2019; Hondzinski et al., 2016; Poirier et al., 2022, 2020). More, recent works confirm kinematics results by revealing systematic negative epochs in the phasic activity of antigravity muscles (Gaveau et al., 2021; Poirier et al., 2022). Such negativity of the phasic signal demonstrate that gravity effects are not compensated for but, instead, harvested to discount muscle effort (Gaveau et al. 2021). These kinematics and electromyographic results are thought to represent the hallmarks of a gravity-related optimization processes that occurs during motor planning to minimize muscle effort (for reviews see (Berret et al., 2019; White et al., 2020). In our recent study (Poirier et al., 2022), we found that direction-dependent kinematics and negative epochs were still present during movements performed with the non-dominant arm, but significantly reduced compared to movements performed with the dominant arm. These results support the hypothesized superiority of the dominant motor system for open-loop optimization processes.

A prominent theory regarding the aging brain is the reduction of hemispheric asymmetries in older adults (HAROLD model; Cabeza, 2002). Neurophysiological studies have reported greater bilateral pattern of activation in older adults during cognitive (Cabeza et al., 2002) and motor tasks (Seidler et al., 2010). Moreover, an age-related reduction of motor lateralization has been shown for actual (Przybyla et al., 2011) and mentally simulated (Paizis et al., 2014) arm reaching movements and path tracing (Raw et al., 2011). Here, we compare the lateralization effect on the well-known gravity-related optimal motor control process between young and older adults. Following results suggesting an age-related reduction of motor lateralization and contrary to young adults (Poirier et al., 2022), we hypothesized that gravity-related optimality would be similar between dominant and non-dominant arms in older adults.

## Materials and Methods

### Participants

Forty-one healthy young [27 males and 14 women; mean age = 24 ± 4 (SD) years] and forty-two healthy older volunteers [16 males and 26 women; mean age = 72 ± 6 (SD) years] were included in this study after giving their written informed consent. All participants had normal or corrected-to-normal vision. None present with any neurological or muscular disorders. All participants were right-handed, as attested by a laterality index superior to 60 (Edinburgh Handedness Inventory, Oldfield 1971). A French National ethics committee (2019-A01558-49) approved all procedures. The research was conducted following legal requirements and international norms (Declaration of Helsinki, 1964).

### Experimental protocol

The experimental protocol was similar to previous studies investigating gravity-related arm movement control (Gentili et al., 2007; Le Seac’h and McIntyre, 2007; Gaveau and Papaxanthis, 2011a; Gaveau et al., 2014, 2016; Poirier et al., 2020). We asked participants to perform unilateral single degree-of-freedom vertical arm movements (rotation around the shoulder joint) in a parasagittal plane (Fig.1A). In a randomized block order, movements were executed with the dominant (D) and non-dominant (ND) arms (Poirier et al., 2022). We investigated single degree-of-freedom movements to isolate and emphasize the mechanical effects of gravity on arm motion. During these movements, the work of gravity torque varies with movement direction, whereas inertia remains constant. To maximally vary the effect of gravity torque on the movement, participants moved in two opposed directions: upward and downward. Directions were randomly interleaved within each block. Each participant carried out 64 trials (16 trials x 2 directions x 2 arms).

**Figure 1.**
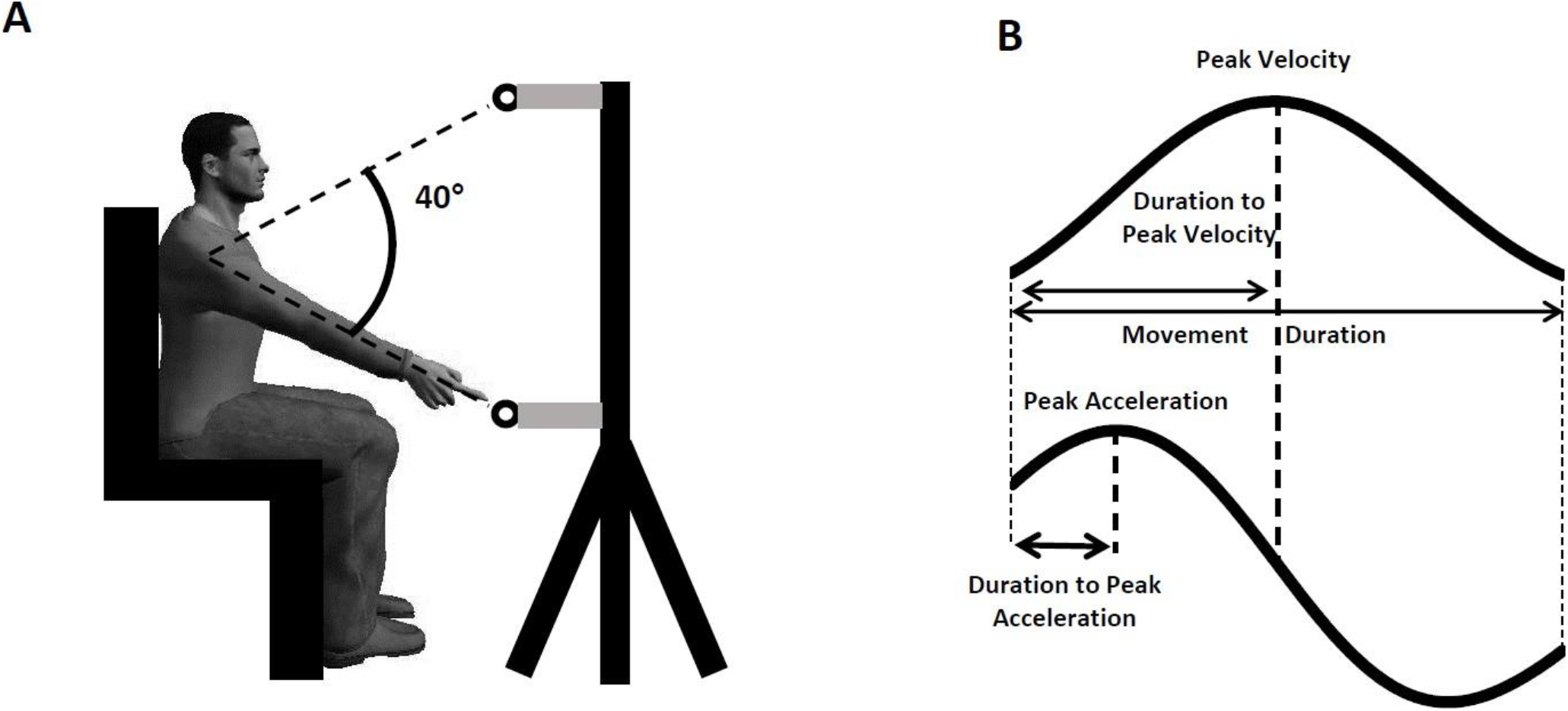
(A) Experimental setup. Participants performed arm pointing movements between two targets on separated trials. (B) Illustration of the parameters computed on the velocity and acceleration profiles.

Participants sat on a chair with their trunk parallel to the gravity vector. Two targets (diameter = 3 cm) were positioned in the parasagittal plane crossing the participant’s right or left shoulder. Targets were placed at a distance corresponding to the length of the participant’s fully extended arm plus two centimeters. Vertical positions of targets were tuned for each participant to require a 40° shoulder flexion or extension. Targets were centered around the antero-posterior horizontal line crossing the shoulder joint, thereby corresponding to a 110° (upward target) and 70° (downward target) shoulder elevation.

A trial was carried out as follows: the participant positioned her/his fully extended arm in front of the initial target (verbally indicated by the experimenter using a color code). After a brief delay (∼2 seconds), the experimenter informed the participant that she/he was free to reach to the other target, as fast and accurately as possible, whenever she/he wanted. The participant had been informed that the reaction time was not constrained. At the end of a movement, participants were requested to maintain their final position (about 2 seconds) until the experimenter instructed them that they were free to relax their arm. To prevent muscle fatigue, participants were invited to have a rest between trials (∼10 s) and blocks (5mn). Before each block, participants were invited to perform few practice trials (∼5 trials) to familiarize with the task.

We placed five reflective markers on the participant’s shoulder (acromion), arm (middle of the humeral bone), elbow (lateral epicondyle), wrist (cubitus styloid process), and finger (nail of the index). We also placed markers on both targets. Three-dimensional position was recorded using an optoelectronic motion capture system (Vicon system, Oxford, UK; six cameras; 100Hz). After calibration, spatial variable error of the system was less than 0.5mm. Additionally, we recorded muscle activation patterns by means of bipolar surface electrodes positioned over the anterior (DA) and posterior (DP) heads of deltoid muscles (Aurion, Zerowire EMG, sampling frequency: 1000Hz). The Giganet unit (Vicon, Oxford, UK) allowed synchronized recording of kinematic and EMG signals.

### Data analysis

We processed Kinematic and EMG data using custom programs written in Matlab (Mathworks, Natick, NA). Data processing was similar to previous studies (Poirier et al., 2020, 2022; Gaveau et al., 2021).

#### Kinematics

We filtered position (third-order low-pass Butterworth filter; 5Hz cut-off, zero-phase distortion, “butter” and “filtfilt” functions) before computing velocity and acceleration profiles by numerical differentiation (3 points derivative). We computed angular joint displacements to ensure that elbow and wrist rotations were negligible (<1° for each trial). Movement onset and offset were respectively defined as the times when finger velocity rose above and fall below a 10% of peak velocity threshold. Movements presenting more than one local maxima, i.e. more than one velocity peak, were automatically rejected from further analysis. The following parameters were computed to quantify kinematic patterns (Fig.1B): 1) Movement duration (MD = offset - onset). 2) Movement amplitude (Amp = offset angle – onset angle). 3) Relative duration to peak acceleration (rD-PA = duration to peak acceleration divided by MD). 4) Relative duration to peak velocity (rD-PV = duration to peak velocity divided by MD).

#### Electromyography

We filtered (bandpass third-order Butterworth filter; 20-300Hz, zero-phase distortion, “butter” and “filtfilt” functions) and then rectified EMG signals before integrating them using a 50ms sliding window from 250ms before movement onset to 250ms after movement offset. EMG signal were then normalized by the EMG recorded during a maximal voluntary isometric contraction (MIVC, same EMG processing as movement patterns). At the beginning of each block, participants performed three MIVC in both flexions and extensions directions, at a 90° shoulder angle. Lastly, we ordered EMG traces according to movement mean velocity and averaged them across two trials (from the two slowest to the two fastest movements), resulting in 8 EMG traces to be analyzed for each block and direction. Before averaging two trials together, we normalized the duration of each trial to the mean duration value of these two trials.

To emphasize the role of gravity in the arm motion, we separated the phasic and tonic components of each EMG signal using a well-known subtraction procedure (Flanders and Herrmann, 1992; Buneo et al., 1994; Flanders et al., 1994; D’Avella et al., 2006, 2008; Gaveau et al., 2021; Poirier et al., 2022). We computed averaged values of the integrated phasic EMG signals from 500ms to 250ms before movement onset and from 250 to 500ms after movement offset (Fig.2A). Then, we extracted the theoretical tonic component using a linear interpolation between these two averaged values (Fig.2B). Finally, we computed the phasic component by subtracting the tonic component from the full EMG signal (Fig.2C).

**Figure 2.**
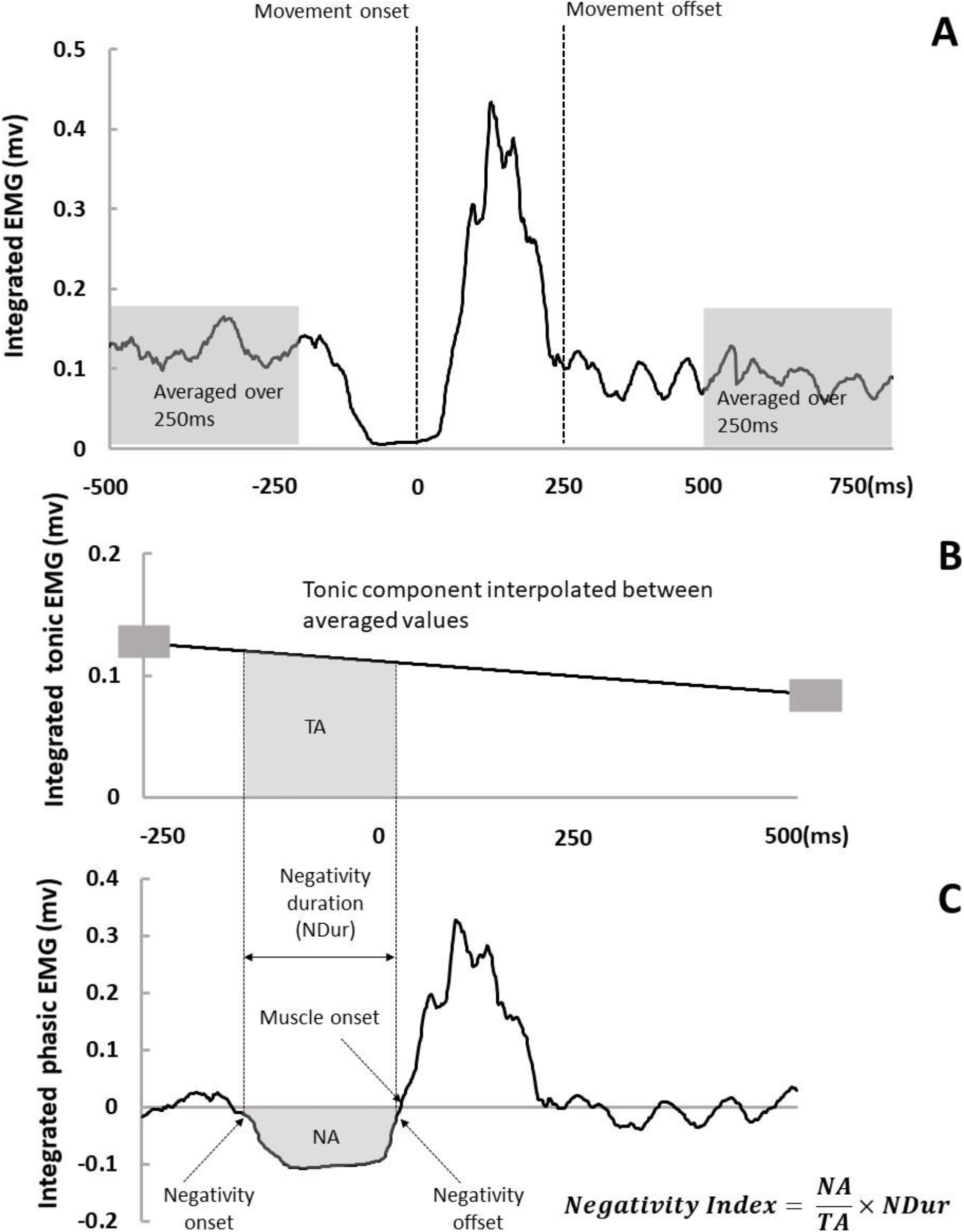
Analysis of the tonic and phasic components of the EMG. (A). Illustration of the integrated full EMG signal. Movement onset and offset were detected on the velocity profile. We averaged the signal from 500 to 250ms before movement onset and from 250 to 500ms after movement offset (grey windows). (B). Tonic EMG component was obtained by linear interpolation between the two previously averaged values (small grey rectangles). (C). Phasic EMG component was obtained by subtracting the tonic component (in panel B) from the full EMG (in panel A). We detected negativity onset and offset and then computed negativity duration (NDur) and integrated phasic signal between negativity onset and offset (NA). TA in panel B represents the integrated tonic signal in the same period.

It was recently shown that phasic EMG activity of antigravity muscles consistently exhibits negative epochs during vertical arm movements (Gaveau et al., 2021) when the arm’s acceleration sign is coherent to gravity’s – i.e. in the acceleration phase of downward movements and in the deceleration phase of upward movements). Model simulations demonstrated that the negativity of antigravity muscles reflects an optimal motor strategy where gravity force is harvested to save muscle force. Recently, we found that the amount of negativity was superior when healthy young participants reach with their dominant than with their non-dominant arm (Poirier et al., 2022). We defined negative epochs as a time interval when phasic EMG was inferior to zero minus a 95% confidence interval (computed on the integrated EMG signals from 500ms to 250ms before movement onset) for at least 50ms. We used this value as a threshold to define negativity onset (when phasic EMG dropped below) and offset (when phasic EMG rose above). Then we computed relative negativity duration (negativity duration divided by movement duration), negativity amplitude (negativity occurrence – i.e. the number of trials including a negative period divided by the total number of trials in the condition – and a negativity index to compare the amount of negativity between conditions:

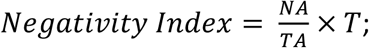

where NA stands for Negative Area and is the integrated phasic signal between negativity onset and offset; TA stands for Tonic Area and is the integrated tonic signal between negativity onset and offset (if NA and TA are equal, it means that the muscle is completely silent during the interval); and T is the duration of the negative epoch normalized by movement duration. We also computed the negativity amplitude, defined as the minimal 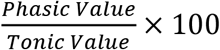 value during the negative phase.

### Statistics

Statistical analyses were performed using SPSS (IBM Corp., Armonk, NY, USA). Distribution normality (Kolgomorov-Smirnov test) and sphericity (Mauchly’s test) was checked for all variables. To analyze movement duration and amplitude, we used repeated measure ANOVA with one between participant factor: *age* (young vs. older) and one within participant factor: *arm* (D vs. ND). As it was demonstrated the movement velocity can influence the gravity-related optimization process signatures (Gaveau and Papaxanthis, 2011), all other parameters were analyzed with a repeated measures ANCOVA in which we set the mean velocity of the movement as a covariate. We used *HSD-Tukey tests* for post-hoc comparisons. The level of significance for all analyses was fixed at *p* = 0.05.

## Results

### Kinematics

All participants performed planar vertical arm movements (shoulder azimuth angle <1° for all trials; n =5312) with single peaked and bell-shaped velocity profiles. A qualitative illustration of the velocity profiles for two typical participants (young and older) is displayed in Fig.3. Building on previous results (Poirier et al., 2022), we aimed to compare directional asymmetries (differences between upward and downward) between arms and groups. Thus, for each kinematic parameter, we computed the directional difference (Down-Up). All kinematic analyses are performed on these directional differences. Nonetheless, values for all kinematic parameters and conditions, including directions, are displayed in Table.1.

**Figure.3.**
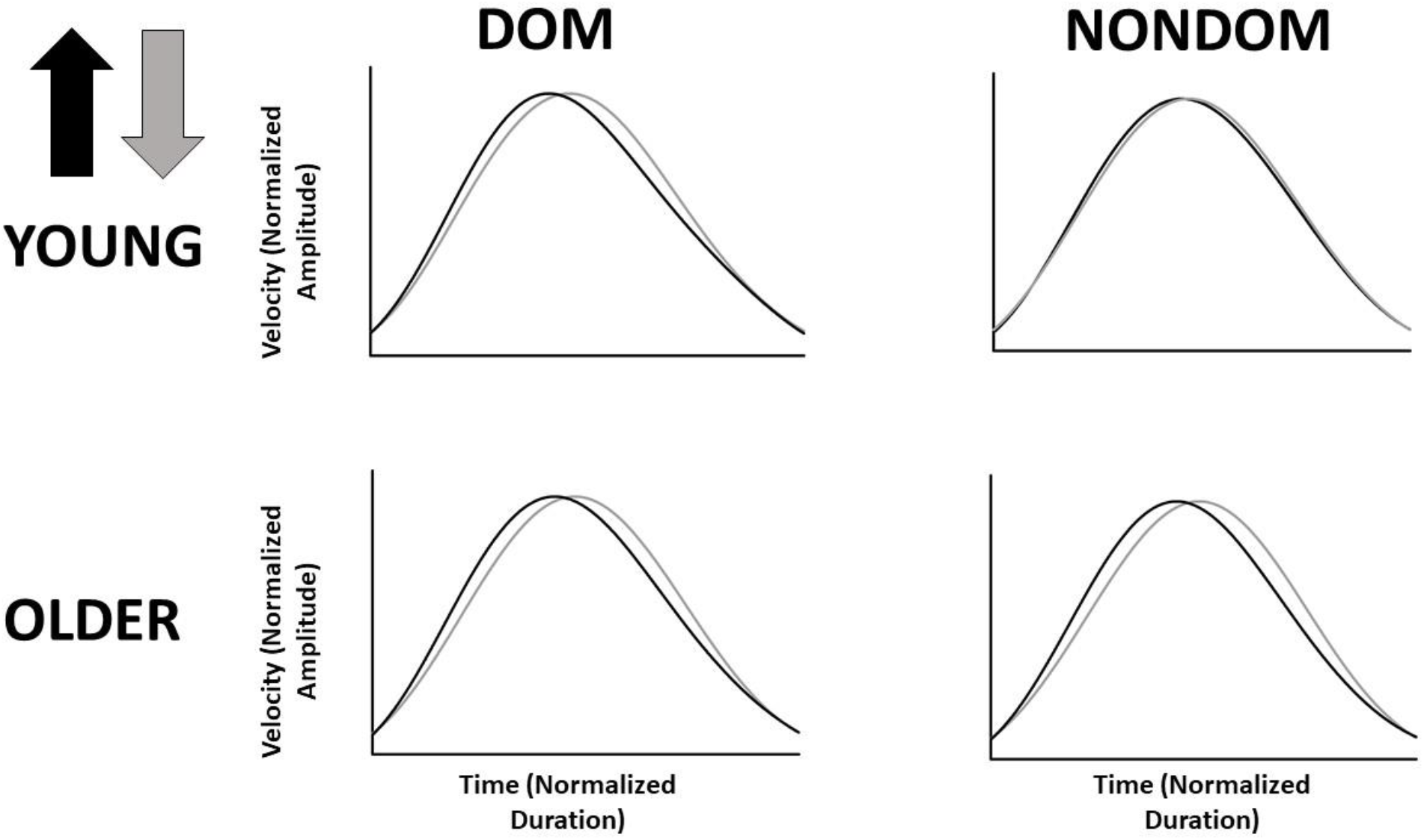
Mean velocity profiles of upward (black traces) and downward (grey traces) movements in a young (upper panels) and an older (lower panels) participants. Left panels represent DOM condition whereas right panels represent NONDOM condition.

**Table.1.**
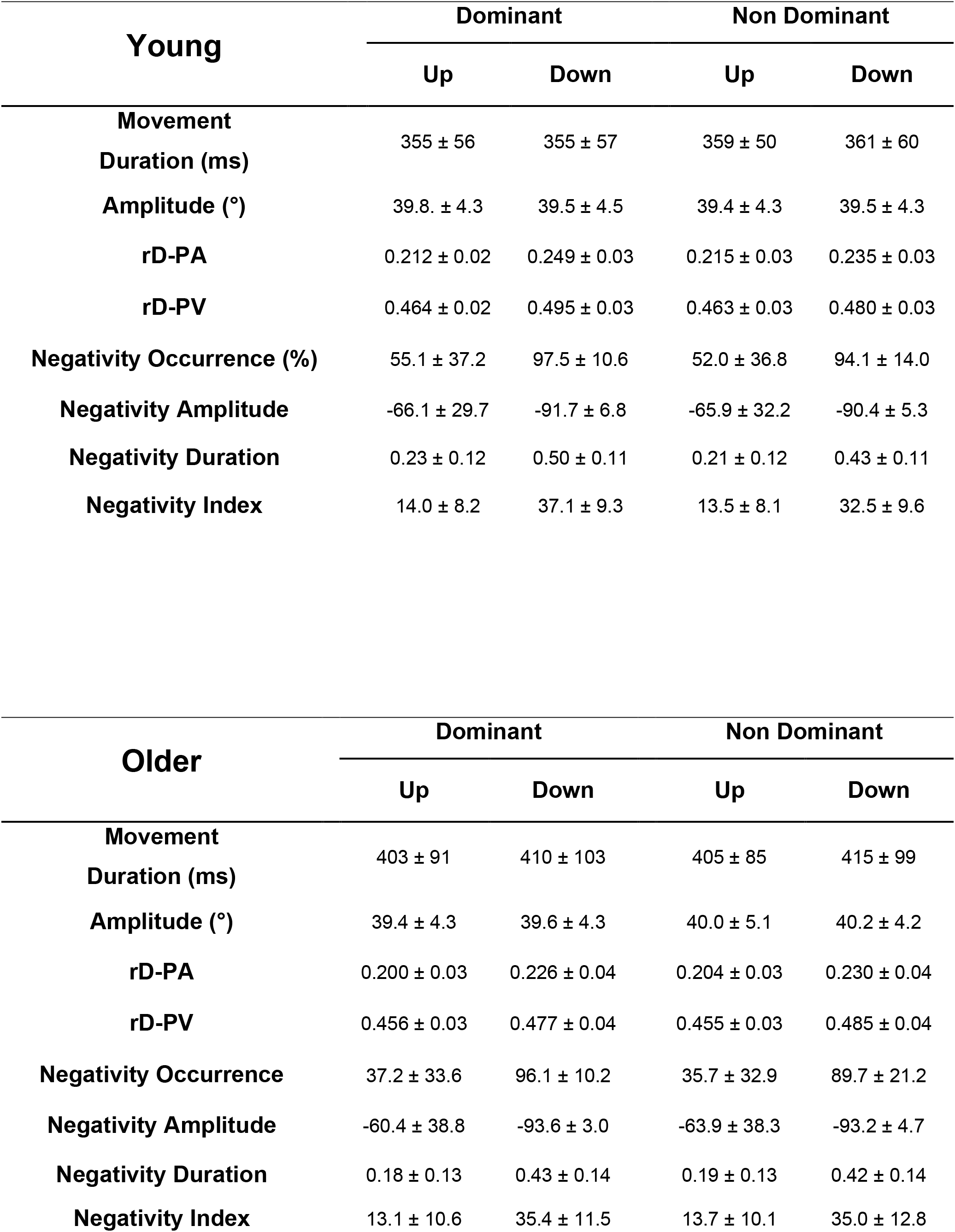
Mean (±SD) values of all kinematic and EMG parameters.

#### Movement Duration and Amplitude

We did not observe any significant effect of *age* (F_(1, 81)_ = 3.50; p = 0.07; η_p_^2^ = 0.04) or *arm* (F_(1, 81)_ = 0.60; p = 0.44; η_p_^2^ = 7e-3) on movement duration. Additionally, *arm x age* interaction effect was not found to be significant (F_(1, 81)_ = 3e-3; p = 0.96; η_p_^2^ = 3e-5). Concerning movement amplitude neither *age* effect (F_(1, 81)_ = 1.11; p = 0.29; η_p_^2^ = 0.01), *arm* (F_(1, 81)_ = 0.50; p = 0.48; η_p_^2^ = 6e-3) nor *arm x age* interaction effect (F_(1, 81)_ = 0.46; p = 0.50; η_p_^2^ = 6e-3) were found to be significant.

#### Relative Duration to Peak Acceleration (rD-PA)

In accordance with multiple previous results, rD-PA was smaller for upward than for downward movements with both arms and for both groups, as attested by positive directional differences (see Fig.4A). Neither *age* (F_(1, 80)_ = 0.57; p = 0.45; η_p_^2^ = 7e-3) nor *arm* (F_(1, 80)_ = 2.89; p = 0.09; η_p_^2^ = 0.04) effect significantly influenced directional differences. However, we observed a significant *arm x age* interaction effect (F_(1, 80)_ = 7.81; p = 6e-3; η_p_^2^ = 0.09) on rD-PA directional differences. As recently reported (Poirier et al., 2022), the directional difference was smaller with the NONDOM than with the DOM arm in young participants (post-hoc comparison: p = 2e-3; Cohen’s d = 0.72). However, no such difference existed between the DOM and NONDOM arms in older participants (p = 0.98; Cohen’s d = 0.08). Additionally, the directional difference on rD-PA neither significantly differed between young and older participants for the DOM (p = 0.10; Cohen’s d = 0.54) nor for the NONDOM (p = 0.69; Cohen’s d = 0.26) arm.

**Figure.4.**
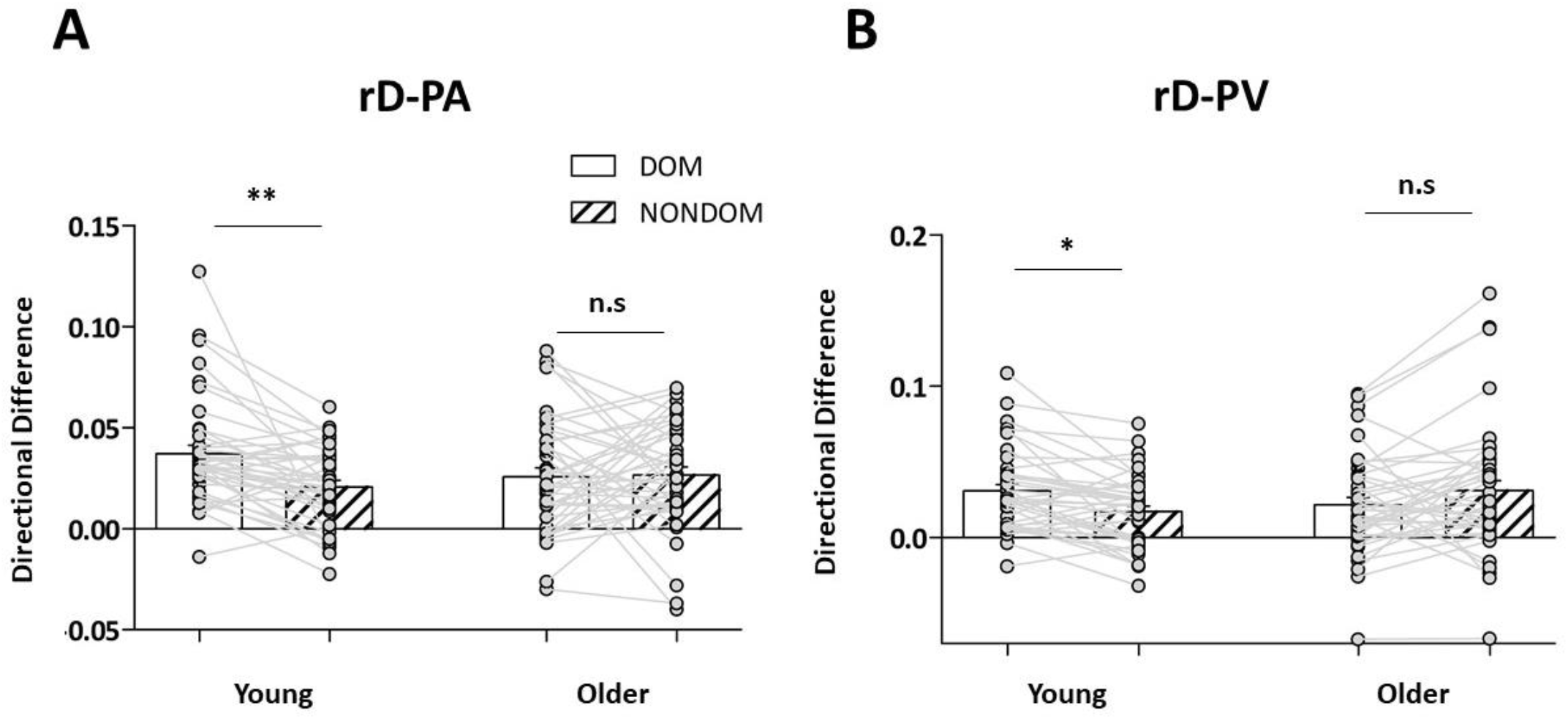
(A). Mean (±SD) values of directional ratios on the relative Duration to Peak Acceleration (rD-PA; computed for each participant as: rD-PA_Down_-rD-PA_Up_). Dominant (DOM) and non-dominant (NONDOM) arm conditions are represented by solid and striped bars, respectively. (B). Mean (±SD) values of directional ratios on the relative Duration to Peak Velocity (rD-PV; computed for each participant as: rD-PV_Down_-rD-PV_Up_). Dominant (DOM) and non-dominant (NONDOM) arm conditions are represented by solid and striped bars, respectively. Grey dots represent individual values.

#### Relative Duration to Peak Velocity (rD-PV)

Here again, rD-PV was smaller for upward than for downward movements with both arms and for both groups, as attested by positive directional differences (see Fig.4B). We did not observe significant main effects of *age* (F_(1, 80)_ = 0.04; p = 0.84; η_p_^2^ = 5e-4) or arm (F_(1, 80)_ = 0.13; p = 0.72; η_p_^2^ = 2e-3). We found a significant *arm x age* interaction effect on rD-PV directional differences (F_(1, 80)_ = 9.51; p = 3e-3; η_p_^2^ = 0.11). Similarly to rD-PA, the directional difference on rD-PV was smaller with the NONDOM than with the DOM arm in young (p = 0.04; Cohen’s d = 0.40) but not in older participants (p = 0.29; Cohen’s d = 0.26). However, there was no significant difference between young and older participants in the DOM (p = 0.39; Cohen’s d = 0.37) nor in the NONDOM (p = 0.61; Cohen’s d = 0.29) conditions.

### EMG

We previously reported systematic negative phases in phasic EMG signal during vertical arm movements (Gaveau et al., 2021; Poirier et al., 2022). These negative phases were shown to occur during the deceleration phase of an upward movement and during the acceleration phase of a downward movement; i.e., when gravity torque helps to produce the arm’s motion. This negativity means that antigravity muscles were less activated than it should have been to compensate for gravity torque. In the present study, negative phases during downward movements were detected in 95% and 93% of movements performed by young and older participants respectively. During upward movements, they were detected in 53% and 37% of movements performed by young and older participants respectively. A qualitative illustration of phasic EMG signal for all muscles and conditions in a typical young and a typical older participant is displayed in Figure.5.

**Figure 5.**
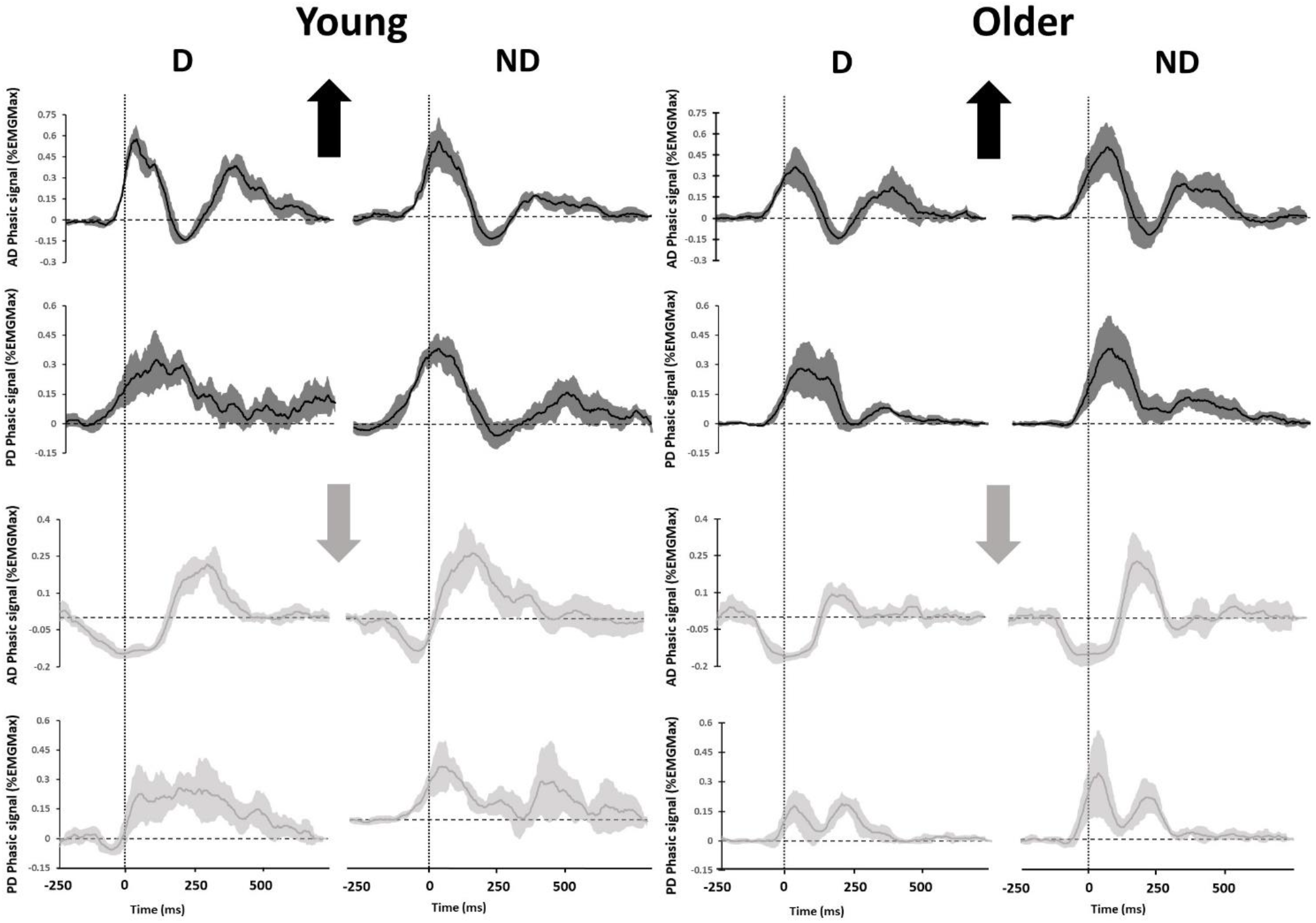
Figure 5. Illustration of the mean (± SD) phasic EMG patterns in a typical young (left panels) and a typical older (right panels) participant for each muscle and condition. Vertical arrows indicate movement directions. Dashed lines represent the kinematic onset of the movement.

In a recent study (Poirier et al., 2022), we also found that the amount of negativity was less important with the NONDOM than with the DOM arm, during downward movements. Similarly to kinematic results in young participants, this result suggests that effort-related motor-optimization is reduced in the NONDOM motor system. Here we aimed to probe optimal motor control lateralization across age. Following previous results, we report statistical analyses on negativity occurrence and index for downward movements. However, we also report the values obtained for negativity amplitude and duration, and for all parameters during upward movements in Table.1. Results for statistical analyses performed on all these parameters are also reported in Table.2.

**Table.2.**
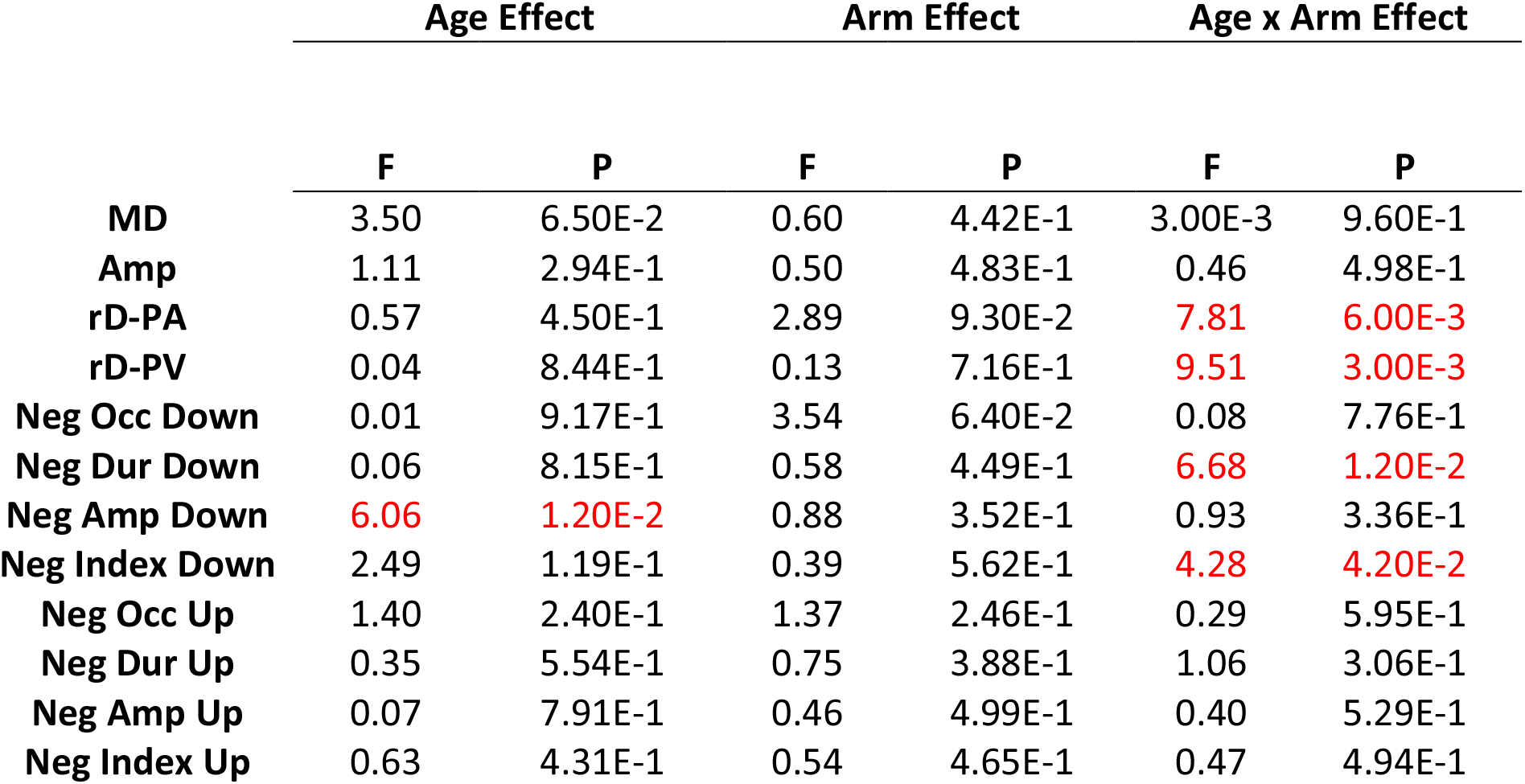
F and P values for all main effects of the ANOVA (MD and Amp) and ANCOVA (all other parameters) are presented for all parameters. Red fonts highlight significant effects. Those statistics are presented for descriptive purposes only. MD: Movement Duration; Amp: Amplitude ; rD-PA; Relative Duration to Peak Acceleration; rD-PV ; Relative Duration to Peak Velocity ; Neg Occ: Negativity Occurrence; Neg Dur: Negativity Duration; Neg Amp; Negativity Amplitude; Neg Index: Negativity Index

|  | Age Effect |  | Arm Effect |  | Age x Arm Effect |  |
| --- | --- | --- | --- | --- | --- | --- |
|  | F | P | F | P | F | P |
| **MD** | 3.50 | 6.50E-2 | 0.60 | 4.42E-1 | 3.00E-3 | 9.60E-1 |
| **Amp** | 1.11 | 2.94E-1 | 0.50 | 4.83E-1 | 0.46 | 4.98E-1 |
| **rD-PA** | 0.57 | 4.50E-1 | 2.89 | 9.30E-2 | **7.81** | **6.00E-3** |
| **rD-PV** | 0.04 | 8.44E-1 | 0.13 | 7.16E-1 | **9.51** | **3.00E-3** |
| **Neg Occ Down** | 0.01 | 9.17E-1 | 3.54 | 6.40E-2 | 0.08 | 7.76E-1 |
| **Neg Dur Down** | 0.06 | 8.15E-1 | 0.58 | 4.49E-1 | **6.68** | **1.20E-2** |
| **Neg Amp Down** | **6.06** | **1.20E-2** | 0.88 | 3.52E-1 | 0.93 | 3.36E-1 |
| **Neg Index Down** | 2.49 | 1.19E-1 | 0.39 | 5.62E-1 | **4.28** | **4.20E-2** |
| **Neg Occ Up** | 1.40 | 2.40E-1 | 1.37 | 2.46E-1 | 0.29 | 5.95E-1 |
| **Neg Dur Up** | 0.35 | 5.54E-1 | 0.75 | 3.88E-1 | 1.06 | 3.06E-1 |
| **Neg Amp Up** | 0.07 | 7.91E-1 | 0.46 | 4.99E-1 | 0.40 | 5.29E-1 |
| **Neg Index Up** | 0.63 | 4.31E-1 | 0.54 | 4.65E-1 | 0.47 | 4.94E-1 |

#### Negativity Occurrence

Negativity occurrence during downward movements was similar between groups and conditions (See Table.1 and Fig.6A). We did not observe significant main effects of *age* (F_(1, 78)_ = 0.01; p = 0.92; η_p_^2^ = 1e-4) nor *arm* (F_(1, 78)_ = 3.54; p = 0.07; η_p_^2^ = 0.04). *Arm x age* interaction effect (F_(1, 78)_ = 0.08; p = 0.77; η_p_^2^ = 1e-3) did not influence negativity occurrence either.

**Figure. 6.**
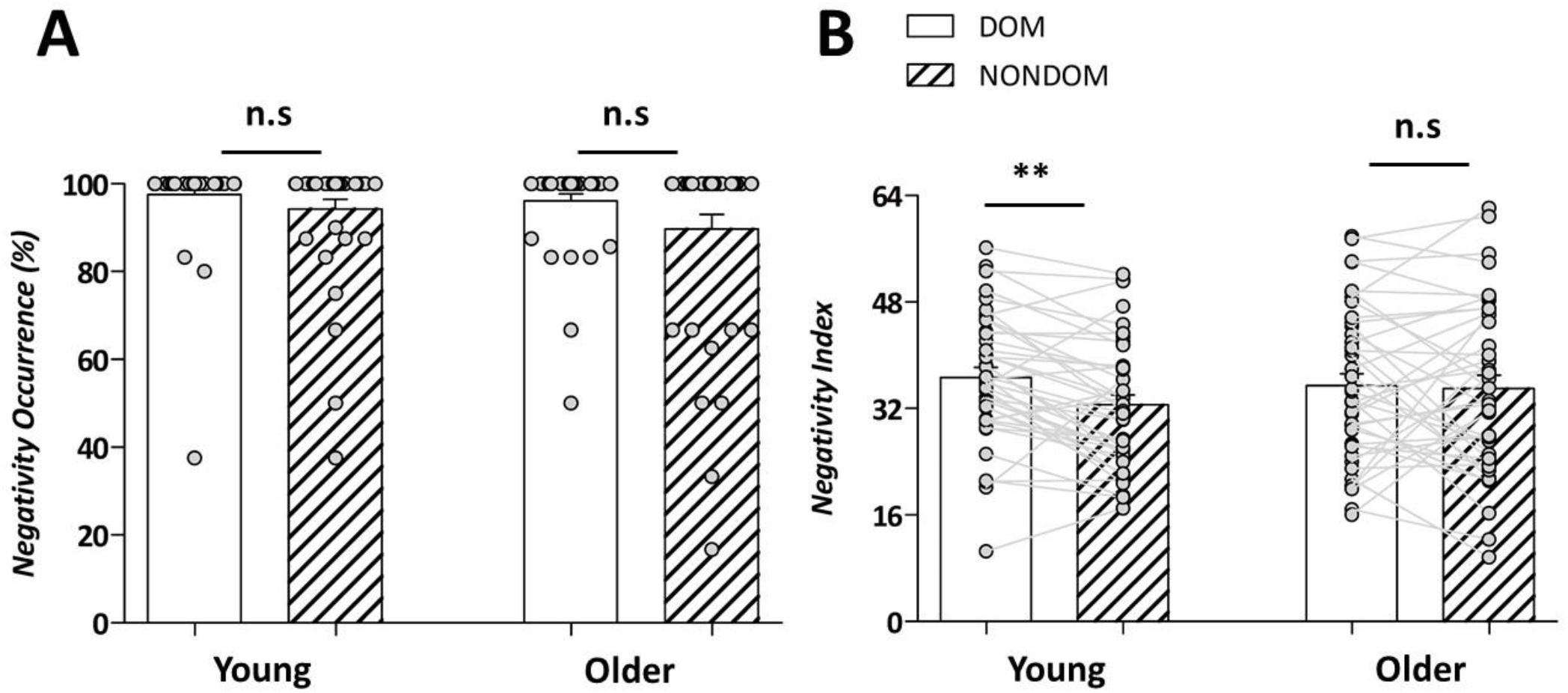
(A). Mean (+SD) negativity occurrence for downward movements. Dominant (DOM) and non-dominant (NONDOM) arm conditions are represented by solid and striped bars, respectively. Grey dots represent individual values. (B). Mean (+SE) negativity index for downward movements. Dominant (DOM) and non-dominant (NONDOM) arm conditions are represented by solid and striped bars, respectively. Grey dots represent individual values.

#### Negativity Index

We also found a significant *arm x age* interaction effect (F_1, 78)_ = 4.28; p = 0.04; η_p_^2^ = 0.05), revealing once again that the difference between arms was greater in young compared to older participants. Post-hoc tests highlighted that negativity index was significantly smaller in NONDOM compared to DOM movements in young (p = 9e-3; Cohen’s d = 0.46; see Fig.6B) but not in older participants (p = 0.99; Cohen’s d = 0.04; see Fig.6B). Here again, negativity index did not differ between young and older participants for the DOM (p = 0.95; Cohen’s d = 0.13) and the NONDOM (p = 0.10; Cohen’s d = 0.55) arms. Main effects of *age* (F_(1, 78)_ = 2.49; p = 0.12; η_p_^2^ = 0.03) and *arm* (F_(1, 78)_ = 0.34; p = 0.56; η_p_^2^ = 4e-3) were not significant.

Altogether, these results demonstrate that the lateralization effect on gravity-related muscle effort minimization decreases with aging.

## Discussion

In this study, we investigated gravity-related optimal motor control in vertical arm movements performed with the dominant and the non-dominant arms in young and older adults. We found that direction-dependent kinematics and negative epochs in phasic muscular activity were reduced for movements performed with the non-dominant arm in young participants. These findings replicate the results of Poirier et al. (2022), confirming the superiority of the dominant motor system in taking advantage of gravity effects to minimize muscle effort in young adults. We found no arm difference in older participants, such that parameters with both arms lied between those of young participants with their dominant and non-dominant arm.

Overall, the present kinematics and EMG results consistently support previous results showing a reduced motor lateralization in older adults. Investigating arm-reaching movements in young and older adults, Przybyla et al. (2011) found interlimb asymmetries concerning movement error and curvature for young but not older participants during reaching movements. In the study of (Raw et al., 2011), young and older participants performed tracing movements with their dominant and their non-dominant hands. They found that young participants were faster with their dominant hand, whereas older participants performed dominant and non-dominant movements with similar speeds. In the study of Paizis et al. (2014), participants mentally simulated and actually executed pointing movements with either their dominant or non-dominant arm in the horizontal plane.

There were inter-limbs asymmetries regarding movement speed (faster movements for the dominant arm) for young participants in both overt and covert movements. In older participants, however, inter-limbs asymmetries were present for overt but not for covert movements. These results are in accordance with the hemispheric asymmetry reduction in older adults (Harold model, Cabeza, 2002) and numerous neurophysiological studies showing more bilateral neural recruitment during motor tasks (Mattay et al., 2002; Heuninckx et al., 2008; Boudrias et al., 2012). From these results, one could hypothesize that, whatever the arm used to perform the movement, older adults recruit motor areas of both hemispheres and therefore produce a less-lateralized motor control.

Additionally, we observed that direction-dependent kinematics are present in older adults and similar to young participants. This result supports the conclusion of a previous study on the fact that optimal motor planning in vertical arm movements is preserved during aging (Poirier et al., 2020). Here we extended these kinematic results to muscle activation patterns by reporting negative epochs on phasic muscular activity (considered as the hallmark of the gravity-related motor optimization process; Gaveau et al., 2021) in older participants. With the dominant arm, the amount of negativity was similar in young and older participants. Thus, the present study provides further evidence for the preservation of motor planning abilities in healthy aging. These abilities are known to depend on predictive processes. Recently, multiple studies suggested that predictive mechanisms remain functional and are increasingly relied upon to compensate for deteriorated sensory cues in aging (Boisgontier and Nougier, 2013; Helsen et al., 2016; Wolpe et al., 2016; Hoellinger et al., 2017; Vandevoorde and Orban de Xivry, 2019). Here we observed an *arm x age* effect on negative epochs of the phasic EMGs during downward movements only. Because negativity appears earlier in downward than in upward movements, this result may support the hypothesized increased reliance on predictive mechanisms in older adults.

Overall, the present results reveal that the diminished motor control lateralization in older adults – that has been largely reported and observed at various levels of investigation, from behavior to brain activations – extends to motor optimization processes. Whether this represents a compensation or a deterioration process remains an open question. Employing behavioral paradigms that target specific motor control processes may help disentangling deterioration from compensation processes in the aging brain (Krakauer et al., 2017; Poirier et al., 2021).

## Bibliography

Bagesteiro, L. B., and Sainburg, R. L. (2002). Handedness: Dominant arm advantages in control of limb dynamics. J. Neurophysiol. 88, 2408–2421. doi:10.1152/jn.00901.2001.

Berret, B., Darlot, C., Jean, F., Pozzo, T., Papaxanthis, C., and Gauthier, J. P. (2008). The inactivation principle: Mathematical solutions minimizing the absolute work and biological implications for the planning of arm movements. PLoS Comput. Biol. 4, e1000194. doi:10.1371/journal.pcbi.1000194.

Berret, B., Delis, I., Gaveau, J., and Jean, F. (2019). “Optimality and modularity in human movement: From optimal control to muscle synergies,” in Springer Tracts in Advanced Robotics, 105–133. doi:10.1007/978-3-319-93870-7_6.

Boisgontier, M. P., and Nougier, V. (2013). Ageing of internal models: From a continuous to an intermittent proprioceptive control of movement. Age (Omaha). 35, 1339–1355. doi:10.1007/s11357-012-9436-4.

Boudrias, M. H., Gonçalves, C. S., Penny, W. D., Park, C. hyun, Rossiter, H. E., Talelli, P., et al. (2012). Age-related changes in causal interactions between cortical motor regions during hand grip. Neuroimage 59, 3398–3405. doi:10.1016/j.neuroimage.2011.11.025.

Buneo, C. A., Soechting, J. F., and Flanders, M. (1994). Muscle activation patterns for reaching: The representation of distance and time. J. Neurophysiol. 71, 1546–1558. doi:10.1152/jn.1994.71.4.1546.

Cabeza, R. (2002). Hemispheric asymmetry reduction in older adults: the HAROLD model. Psychol. Aging 17, 85–100. Available at: http://www.ncbi.nlm.nih.gov/pubmed/11931290 [Accessed August 16, 2019].

Cabeza, R., Anderson, N. D., Locantore, J. K., and McIntosh, A. R. (2002). Aging gracefully: compensatory brain activity in high-performing older adults. Neuroimage 17, 1394–402. Available at: http://www.ncbi.nlm.nih.gov/pubmed/12414279 [Accessed August 16, 2019].

Carson, R. G., Goodman, D., and Elliott, D. (1992). Asymmetries in the discrete and pseudocontinuous regulation of visually guided reaching. Brain Cogn. 18, 169–191. doi:10.1016/0278-2626(92)90077-Y.

Crevecoeur, F., Thonnard, J.-L., and Lefèvre, P. (2009). Optimal integration of gravity in trajectory planning of vertical pointing movements. J. Neurophysiol. 102, 786–96. doi:10.1152/jn.00113.2009.

D’Avella, A., Fernandez, L., Portone, A., and Lacquaniti, F. (2008). Modulation of phasic and tonic muscle synergies with reaching direction and speed. J. Neurophysiol. 100, 1433–1454. doi:10.1152/jn.01377.2007.

D’Avella, A., Portone, A., Fernandez, L., and Lacquaniti, F. (2006). Control of fast-reaching movements by muscle synergy combinations. J. Neurosci. 26, 7791–7810. doi:10.1523/JNEUROSCI.0830-06.2006.

Flanders, M., and Herrmann, U. (1992). Two components of muscle activation: Scaling with the speed of arm movement. J. Neurophysiol. 67, 931–943. doi:10.1152/jn.1992.67.4.931.

Flanders, M., Pellegrini, J. J., and Soechting, J. F. (1994). Spatial/temporal characteristics of a motor pattern for reaching. J. Neurophysiol. 71, 811–813. doi:10.1152/jn.1994.71.2.811.

Flowers, K. (1975). Handedness and controlled movement. Br. J. Psychol. 66, 39–52. doi:10.1111/j.2044-8295.1975.tb01438.x.

Gaveau, J., Berret, B., Angelaki, D. E., and Papaxanthis, C. (2016). Direction-dependent arm kinematics reveal optimal integration of gravity cues. Elife 5. doi:10.7554/eLife.16394.

Gaveau, J., Berret, B., Demougeot, L., Fadiga, L., Pozzo, T., and Papaxanthis, C. (2014). Energy-related optimal control accounts for gravitational load: comparing shoulder, elbow, and wrist rotations. J. Neurophysiol. 111, 4–16. doi:10.1152/jn.01029.2012.

Gaveau, J., Grospretre, S., Berret, B., Angelaki, D. E., and Papaxanthis, C. (2021). A cross-species neural integration of gravity for motor optimization. Sci. Adv. 7, eabf7800. doi:10.1126/sciadv.abf7800.

Gaveau, J., Paizis, C., Berret, B., Pozzo, T., and Papaxanthis, C. (2011). Sensorimotor adaptation of point-to-point arm movements after spaceflight: the role of internal representation of gravity force in trajectory planning. J. Neurophysiol. 106, 620–9. doi:10.1152/jn.00081.2011.

Gaveau, J., and Papaxanthis, C. (2011a). The temporal structure of vertical arm movements. PLoS One 6, e22045. doi:10.1371/journal.pone.0022045.

Gaveau, J., and Papaxanthis, C. (2011b). The Temporal Structure of Vertical Arm Movements. PLoS One 6, e22045. doi:10.1371/journal.pone.0022045.

Gentili, R., Cahouet, V., and Papaxanthis, C. (2007). Motor planning of arm movements is direction-dependent in the gravity field. Neuroscience 145, 20–32. doi:10.1016/j.neuroscience.2006.11.035.

Helsen, W. F., Van Halewyck, F., Levin, O., Boisgontier, M. P., Lavrysen, A., and Elliott, D. (2016). Manual aiming in healthy aging: does proprioceptive acuity make the difference? Age (Omaha). 38, 45. doi:10.1007/s11357-016-9908-z.

Heuninckx, S., Wenderoth, N., and Swinnen, S. P. (2008). Systems neuroplasticity in the aging brain: Recruiting additional neural resources for successful motor performance in elderly persons. J. Neurosci. 28, 91–99. doi:10.1523/JNEUROSCI.3300-07.2008.

Hoellinger, T., McIntyre, J., Jami, L., Hanneton, S., Cheron, G., and Roby-Brami, A. (2017). A strategy of faster movements used by elderly humans to lift objects of increasing weight in ecological context. Neuroscience 357, 384–399. doi:10.1016/j.neuroscience.2017.04.010.

Hondzinski, J. M., Soebbing, C. M., French, A. E., and Winges, S. A. (2016). Different damping responses explain vertical endpoint error differences between visual conditions. Exp. Brain Res. 234, 1575–1587. doi:10.1007/s00221-015-4546-8.

Jayasinghe, S. AL, Sarlegna, F. R., Scheidt, R. A., and Sainburg, R. L. (2021). Somatosensory deafferentation reveals lateralized roles of proprioception in feedback and adaptive feedforward control of movement and posture. Curr. Opin. Physiol. 19, 141–147. doi:10.1016/j.cophys.2020.10.005.

Krakauer, J. W., Ghazanfar, A. A., Gomez-Marin, A., MacIver, M. A., and Poeppel, D. (2017). Neuroscience Needs Behavior: Correcting a Reductionist Bias. Neuron 93, 480–490. doi:10.1016/j.neuron.2016.12.041.

Le Seac’h, A. B., and McIntyre, J. (2007). Multimodal reference frame for the planning of vertical arms movements. Neurosci. Lett. 423, 211–215. doi:10.1016/j.neulet.2007.07.034.

Mattay, V. S., Fera, F., Tessitore, A., Hariri, A. R., Das, S., Callicott, J. H., et al. (2002). Neurophysiological correlates of age-related changes in human motor function. Neurology 58, 630–635. doi:10.1212/WNL.58.4.630.

Maurus, P., Kurtzer, I., Antonawich, R., and Cluff, T. (2021). Similar stretch reflexes and behavioral patterns are expressed by the dominant and nondominant arms during postural control. https://doi.org/10.1152/jn.00152.2021 126, 743–762. doi:10.1152/JN.00152.2021.

Oldfield, R. C. (1971). The assessment and analysis of handedness: The Edinburgh inventory. Neuropsychologia 9, 97–113. doi:10.1016/0028-3932(71)90067-4.

Paizis, C., Skoura, X., Personnier, P., and Papaxanthis, C. (2014). Motor asymmetry attenuation in older adults during imagined arm movements. Front. Aging Neurosci. 6, 49. doi:10.3389/fnagi.2014.00049.

Papaxanthis, C., Pozzo, T., and McIntyre, J. (2005). Kinematic and dynamic processes for the control of pointing movements in humans revealed by short-term exposure to microgravity. Neuroscience 135, 371–383. doi:10.1016/j.neuroscience.2005.06.063.

Papegaaij, S., Taube, W., Baudry, S., Otten, E., and Hortobágyi, T. (2014). Aging causes a reorganization of cortical and spinal control of posture. Front. Aging Neurosci. 6, 28. doi:10.3389/fnagi.2014.00028.

Poirier, G., Ohayon, A., Juranville, A., Mourey, F., and Gaveau, J. (2021). Deterioration, Compensation and Motor Control Processes in Healthy Aging, Mild Cognitive Impairment and Alzheimer’s Disease. Geriatrics 6, 33. doi:10.3390/geriatrics6010033.

Poirier, G., Papaxanthis, C., Mourey, F., and Gaveau, J. (2020). Motor Planning of Vertical Arm Movements in Healthy Older Adults: Does Effort Minimization Persist With Aging? Front. Aging Neurosci. 12. doi:10.3389/fnagi.2020.00037.

Poirier, G., Papaxanthis, C., Mourey, F., Lebigre, M., and Gaveau, J. (2022). Muscle effort is best minimized by the right-dominant arm in the gravity field. J. Neurophysiol. 127, 1117–1126. doi:10.1152/jn.00324.2021.

Przybyla, A., Haaland, K. Y., Bagesteiro, L. B., and Sainburg, R. L. (2011). Motor asymmetry reduction in older adults. Neurosci. Lett. 489, 99–104. doi:10.1016/j.neulet.2010.11.074.

Raw, R. K., Wilkie, R. M., Culmer, P. R., and Mon-Williams, M. (2011). Reduced motor asymmetry in older adults when manually tracing paths. Exp. Brain Res. 2011 2171 217, 35–41. doi:10.1007/S00221-011-2971-X.

Roy, E. A., and Elliott, D. (1986). Manual asymmetries in visually directed aiming. Can. J. Psychol. 40, 109–121. doi:10.1037/h0080087.

Roy, E. A., Kalbfleisch, L., and Elliott, D. (1994). Kinematic analyses of manual asymmetries in visual aiming movements. Brain Cogn. 24, 289–295. doi:10.1006/brcg.1994.1017.

Sainburg, R. L. (2002). Evidence for a dynamic-dominance hypothesis of handedness. Exp. Brain Res. 142, 241–258. doi:10.1007/s00221-001-0913-8.

Sainburg, R. L. (2014). Convergent models of handedness and brain lateralization. Front. Psychol. 5, 1092. doi:10.3389/fpsyg.2014.01092.

Schaffer, J. E., and Sainburg, R. L. (2017). Interlimb differences in coordination of unsupported reaching movements. Neuroscience 350, 54–64. doi:10.1016/J.NEUROSCIENCE.2017.03.025.

Seidler, R. D., Bernard, J. A., Burutolu, T. B., Fling, B. W., Gordon, M. T., Gwin, J. T., et al. (2010). Motor control and aging: links to age-related brain structural, functional, and biochemical effects. Neurosci Biobehav Rev 34, 721–733. doi:10.1016/j.neubiorev.2009.10.005.

Takagi, A., Maxwell, S., Melendez-Calderon, A., and Burdet, E. (2020). The dominant limb preferentially stabilizes posture in a bimanual task with physical coupling. https://doi.org/10.1152/jn.00047.2020 123, 2154–2160. doi:10.1152/JN.00047.2020.

Vandevoorde, K., and Orban de Xivry, J. J. (2019). Internal model recalibration does not deteriorate with age while motor adaptation does. Neurobiol. Aging 80, 138–153. doi:10.1016/j.neurobiolaging.2019.03.020.

White, O., Gaveau, J., Bringoux, L., and Crevecoeur, F. (2020). The gravitational imprint on sensorimotor planning and control. J. Neurophysiol. doi:10.1152/jn.00381.2019.

Wolpe, N., Ingram, J. N., Tsvetanov, K. A., Geerligs, L., Kievit, R. A., Henson, R. N., et al. (2016). Ageing increases reliance on sensorimotor prediction through structural and functional differences in frontostriatal circuits. Nat. Commun. 7. doi:10.1038/ncomms13034.

Woytowicz, E. J., Westlake, K. P., Whitall, J., and Sainburg, R. L. (2018). Handedness results from complementary hemispheric dominance, not global hemispheric dominance: evidence from mechanically coupled bilateral movements. J. Neurophysiol. 120, 729–740. doi:10.1152/jn.00878.2017.

Yadav, V., and Sainburg, R. L. (2011). Motor lateralization is characterized by a serial hybrid control scheme. Neuroscience 196, 153–167. doi:10.1016/j.neuroscience.2011.08.039.

Yadav, V., and Sainburg, R. L. (2014). Limb dominance results from asymmetries in predictive and impedance control mechanisms. PLoS One 9, e93892. doi:10.1371/journal.pone.0093892.

Yamamoto, S., Fujii, K., Zippo, K., Kushiro, K., and Araki, M. (2019). The kinetic mechanisms of vertical pointing movements. Heliyon 5, e02012. doi:10.1016/J.HELIYON.2019.E02012.

Yamamoto, S., and Kushiro, K. (2014). Direction-dependent differences in temporal kinematics for vertical prehension movements. Exp. Brain Res. 232, 703–711. doi:10.1007/s00221-013-3783-y.

Yamamoto, S., Shiraki, Y., Uehara, S., and Kushiro, K. (2016). Motor control of downward object-transport movements with precision grip by object weight. Somatosens. Mot. Res. 33, 130–136. doi:10.1080/08990220.2016.1203304.

